# Prediction of pathological subthalamic nucleus beta burst occurrence in Parkinson’s disease

**DOI:** 10.1101/2024.05.09.593398

**Authors:** Bahman Abdi-Sargezeh, Sepehr Shirani, Abhinav Sharma, Alexander Green, Harith Akram, Ludvic Zrinzo, Patricia Limousin, Tom Foltynie, Timothy Denison, Huiling Tan, Vladimir Litvak, Ashwini Oswal

**Affiliations:** MRC Brain Network Dynamics Unit, University of Oxford, Oxford, UK; Nuffield Department of Clinical Neurosciences, University of Oxford, Oxford, UK; Institute of Psychiatry, Psychology & Neuroscience, Department of Basic and Clinical Neuroscience, King’s College London, London, UK; Department of Clinical and Movement Neurosciences, University College London, London, UK; Wellcome Centre for Human Neuroimaging, University College London, UK

**Keywords:** Beta burst prediction, DBS, deep learning, adaptive deep brain stimulation, local field potential

## Abstract

The cortico-basal ganglia network in Parkinson’s disease (PD) is characterized by the emergence of transient episodes of exaggerated beta frequency oscillatory synchrony known as bursts. Although it is well established that bursts of prolonged duration associate closely with motor impairments, the mechanisms leading to burst initiation remain poorly understood. Crucially, it is unclear whether there are features of basal ganglia activity which reliably predict burst onset. Current adaptive Deep Brain Stimulation (aDBS) approaches can only reactively deliver stimulation following burst detection and are unable to stimulate proactively to prevent burst onset. The discovery of predictive biomarkers could allow for such proactive stimulation, thereby offering potential for improvements in therapeutic efficacy. Here, using deep learning, we show that the timing of subthalamic nucleus (STN) beta bursts can be accurately predicted up to 60 ms prior to onset. Furthermore, we highlight that a dip in the beta amplitude - which is likely to be indicative of a phase reset of oscillatory populations occurring between 80-100 ms prior to burst onset - is a predictive biomarker for burst occurrence. These findings demonstrate proof-of-principle for the feasibility of beta burst prediction for DBS and provide insights into the mechanisms of burst initiation.

## Introduction

Parkinson’s disease (PD) is a common neurodegenerative condition which is characterised by nigrostriatal dopamine depletion and the emergence of stereotyped patterns of oscillatory synchrony within cortico-basal ganglia circuits^1,2^. Excessive synchronisation across the beta frequency range (13-30 Hz) characterises the parkinsonian dopamine depleted state and is believed to relate directly to motoric impairment^3,4^. Therapeutic approaches such as both STN DBS and dopaminergic medication lead to a suppression of basal ganglia beta oscillatory synchrony, with the degree of suppression correlating positively with motor improvements^5–8^. Furthermore, a causal effect of beta oscillations on movement is suggested by the observation that entraining motor cortical beta rhythms results in movement slowing^9^.

Recent observations highlight that beta activity is not continuous but occurs in short-lived packets known as **bursts** ^10^. Although the mechanisms of beta burst generation remain poorly understood, it is increasingly believed that **bursts** of longer duration and amplitude may be particularly detrimental to motor function in PD^11^. This finding has led to beta activity being used as a control signal in amplitude-responsive closed loop DBS, where stimulation is delivered only when beta amplitude rises above a certain threshold^12–16^. Studies reveal that beta triggered adaptive DBS (aDBS) is more effective than conventional continuous DBS^15,16^. Additionally, by virtue of selectively targeting a pathophysiological signal of interest, aDBS may offer additional benefits including reduced stimulation requirement and a lower incidence of stimulation induced side effects such as dyskinesia, gait impairment and speech impairment^17^.

One drawback of aDBS is that stimulation is initiated after some fixed delay following the actual occurrence of a **burst**^15^. This delay, which can be up to hundreds of milliseconds, will be the sum of the time taken for the burst to be detected and the system delay between burst detection and stimulation initiation^15^. This means that aDBS can only reactively suppress beta bursts after they have developed and propagated within the cortico-basal ganglia circuit. The discovery of a reliable biomarker that could allow for the **prediction** of bursts would facilitate the development of **proactive DBS** approaches with the capability of either preventing bursts or suppressing them earlier following their onset^18^. Proactive DBS approaches are likely to be more energy efficient ^19^ and could lead to improvements in both the efficacy and the side effect profile of DBS for PD.

In this work, we test the **hypothesis** that STN activity includes features which reliably **predict** beta **burst** onset. To address this question, we developed a deep neural network architecture - based on Convolutional Neural Networks (**CNNs**) - that modelled burst prediction as a binary **classification** problem (outputting either 1 or 0 depending on whether or not a burst was expected to occur). Our network was trained on STN local field potential (LFP) activity recorded from PD patients undergoing functional neurosurgery for the insertion of STN DBS electrodes. Importantly, we train and test our neural network architecture on patient specific beta band filtered STN activity, which has emerged as a robust biomarker of motoric state for aDBS applications ^15,20–22^.

Our trained network was able to reliably **predict** burst onset in unseen test data. Furthermore, by considering predictive segments within the test data, we reveal a pathophysiological **state transition** characterised by a reduction followed by a rise in the beta amplitude, which occurs prior to the onset of each burst. Our findings provide proof of principle for the feasibility of **burst prediction** for **proactive** DBS, as well as shedding light on the pathophysiological mechanisms of beta burst initiation.

## Methods

### Patients and experimental details

We studied STN activity in 16 PD patients undergoing bilateral implantation of STN DBS electrodes at the National Hospital for Neurology and Neurosurgery (UCL). In all cases, a Medtronic model 3389 electrode with four platinum-iridium contacts was implanted. Recordings were performed 3-6 days after electrode implantation, before connection and insertion of the implantable pulse generator (see Error! Reference source not found. for further clinical details). Further details of the surgical procedure can be found in other reports ^23,24^. PD diagnoses were made in accordance with the Queen Square Brain Bank Criteria^25^. All patients provided written informed consent and research protocols were approved by the local research ethics committee.

To maximise the probability of beta burst occurrence, recordings were performed following overnight withdrawal from dopaminergic medication (OFF medication). LFP activity was collected using a battery powered and mains optically isolated BrainAmp system (Brain Products) with a sampling frequency of 2400 Hz. Three bipolar channels (0-1, 1-2, 2-3) were recorded from each electrode and the data were subsequently high pass filtered at 1 Hz in the hardware to avoid amplifier saturation due to large DC offsets. Although recordings of cortical activity using magnetoencephalography (MEG) ^5,18,26,27^ were collected at the same time as LFP recordings, in this analysis we focus explicitly on beta burst prediction from the STN LFP alone.

A neurologist was present during the recordings and patients were requested to keep their eyes open and to remain still. Either one or two rest recording sessions were performed. The duration of each session varied between 188 and 253 seconds (with a mean and standard error of the mean (SEM) of 198±4, see Error! Reference source not found.).

### Determination of beta peak frequency and annotation of beta bursts

To determine the beta peak frequency, the power spectrum of the STN LFP from each bipolar contact was obtained using the short-time Fourier transform (STFT). A Hamming window with a length of 1 second and an overlap of 50% was used for spectral estimation. The squared magnitude of the resulting complex spectrum was computed and averaged across all time windows for visualisation between frequencies of 1 and 100 Hz (with a resolution of 1 Hz; see Error! Reference source not found.**a** for an exemplar spectrum). For each participant, the single bipolar LFP channel from each hemisphere with the highest amplitude peak within the beta frequency range (13-30 Hz) - which we term the **beta channel** - was selected for further analysis.

LFP time series from the beta channel were bandpass filtered within a ±3 Hz window centred on the beta peak frequency, using a causal 6th order Butterworth filter implemented in the SciPy library for Python (https://scipy.org/). The application of a causal filter served to prevent future samples from impacting current or past filter outputs which would be used for prediction. We limited our analysis to individual patient beta band filtered signals as these are typically used in aDBS implementations^15,20,21^. After filtering, data were downsampled to 600 Hz (to reduce computational cost) and rectified prior to the peak values being linearly interpolated to produce the beta amplitude envelope of the signal. Finally, beta burst timings were defined as time points where the beta amplitude envelope exceeded its 75th percentile^10,20^ (see Figure 1b). The number of bursts, their duration, and the number of bursts per second are illustrated for each hemisphere separately in Table 1. The mean duration of bursts was 136.4 and 143.6 ms for right and left hemispheres. The mean number of bursts per second was 1.4 for both hemispheres.

**Table 1.** Clinical characteristics of patients and details of STN recordings. LDE = levodopa dose equivalent. The total pre-operative UPDRS part III score is presented in the on and off medication states. S1 and S2 indicate duration of STN recordings for sessions 1 and 2. Burst characteristics are presented separately for data from the right (R) and left (L) hemispheres.

| Case | Age and gender | Disease duration (years) | Preoperative medication (mg) | UPDRS III pre-op off/on | Data length (s) S1/ S2 | No. of bursts R/L | Mean duration of bursts (ms) R/L | No. of bursts per second R/L |
| --- | --- | --- | --- | --- | --- | --- | --- | --- |
| 1 | 54 F | 10 | LDE 1158 | 35/9 | 192/193 | 667/623 | 126.6/123.3 | 1.7/1.6 |
| 2 | 54 M | 15 | LDE 1150 | 53/19 | 190/193 | 545/509 | 148.3/173.3 | 1.4/1.3 |
| 3 | 58 F | 3 | LDE 390 | 41/9 | 191 | 305/262 | 133.3/144.1 | 1.6/1.4 |
| 4 | 47 M | 8 | LDE 2264 | 46/4 | 190/188 | 500/559 | 128.3/126.6 | 1.3/1.5 |
| 5 | 57 M | 8 | LDE 1229 | 41/19 | 238/195 | 513/583 | 130.0/156.6 | 1.2/1.3 |
| 6 | 60 M | 27 | LDE 2048 | 63/8 | 194/189 | 615/597 | 136.6/131.6 | 1.6/1.5 |
| 7 | 57 M | 17 | LDE 1460 | 54/14 | 190 | 279/265 | 138.3/150.0 | 1.4/1.4 |
| 8 | 52 M | 13 | LDE 1484 | 35/10 | 191 | 225/241 | 148.3/151.6 | 1.1/1.3 |
| 9 | 58 M | 11 | LDE 1320 | 43/25 | 201 | 340/304 | 132.5/130.0 | 1.7/1.5 |
| 10 | 72 M | 9 | LDE 1281 | 28/5 | 192 | 316/307 | 127.5/130.0 | 1.6/1.6 |
| 11 | 60 M | 11 | LDE 1012 | 28/5 | 190/201 | 431/675 | 138.3/121.6 | 1.1/1.7 |
| 12 | 43 M | 9 | LDE 1650 | 63/40 | 253 | 346/340 | 134.1/150.8 | 1.3/1.3 |
| 13 | 41 M | 6 | LDE 1220 | 50/22 | 188 | 279/313 | 126.0/128.3 | 1.5/1.7 |
| 14 | 58 M | 12 | LDE 1500 | 38/14 | 202 | 282/250 | 140.8/141.6 | 1.4/1.2 |
| 15 | 60 M | 10 | LDE 1560 | 56/10 | 190 | 205/194 | 175.0/189.2 | 1.1/1.0 |
| 16 | 61 F | 11 | LDE 1049 | 35/4 | 192/225 | 626/566 | 119.1/150.0 | 1.5/1.3 |
| Mean± SEM | 55.7±1.8 | 11.2±1.3 | 1361±107 | 44.3±2.8/<br>13.5±2.4 | 198±4 | 404±38/<br>411±41 | 136.4±3.2/<br>143.6±4.6 | 1.4±0/1.4±0 |

**Figure 1.**
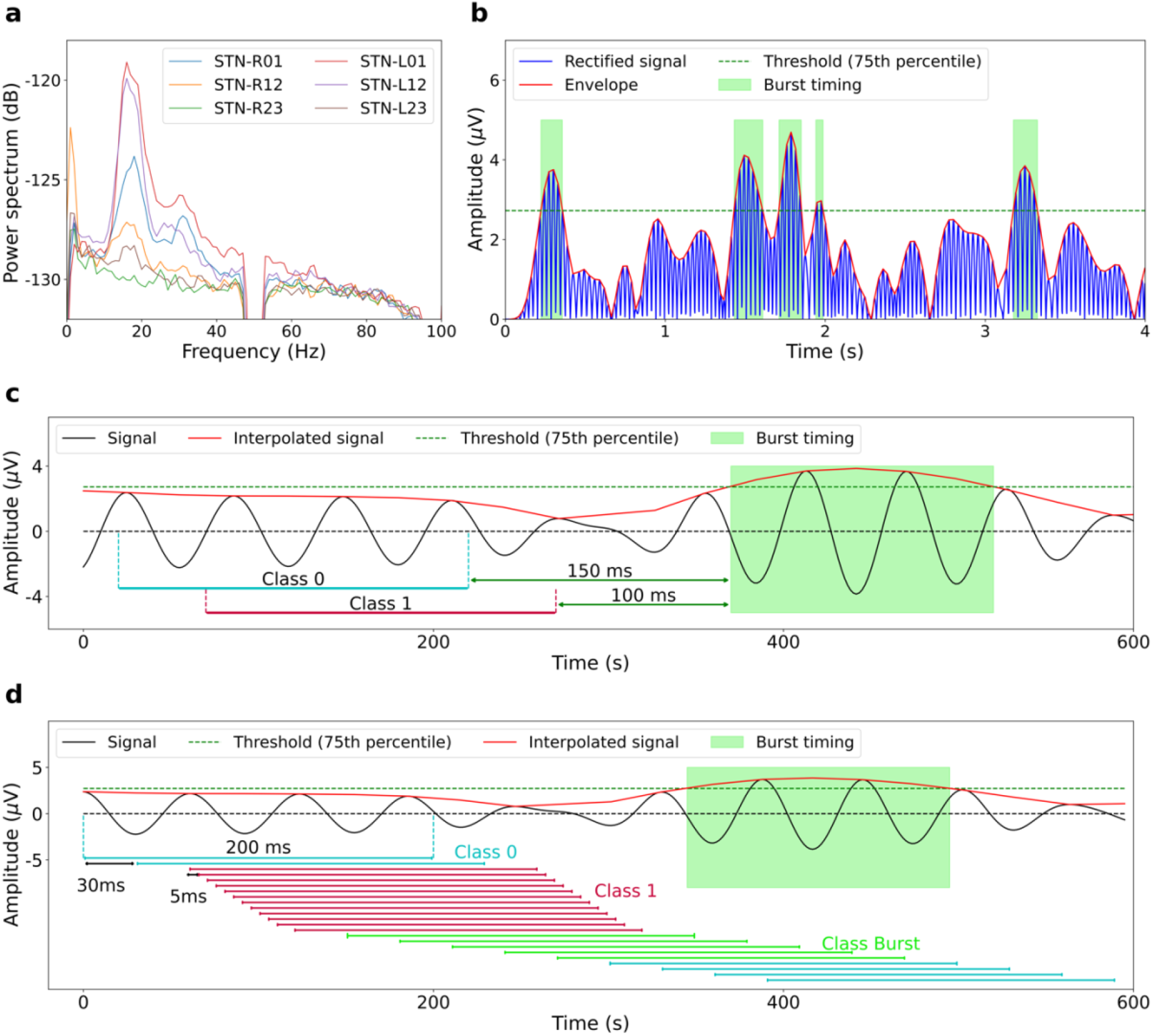
Determination of beta peak frequency, labelling of beta bursts and data segmentation for training of the neural network. (a) Exemplar power spectra of right and left hemisphere STN LFP signals (recorded from bipolar contact pairs 01, 12 and 23) from patient 5 are displayed. In this case, channel STN-R01 from the right hemisphere and channel STN-L01 from the left hemisphere provided the highest amplitude peak within the beta frequency range (peak frequencies at 18 Hz and 16 Hz) and were therefore selected as the *beta channels* for the corresponding hemispheres for further analysis. (b) A 4s long segment of data from the right hemispheric channel STN-R01 of patient 5 is displayed. Data were first filtered (±3 Hz) around the beta peak frequency before being rectified and interpolated. The 75^th^ percentile of the interpolated signal amplitude distribution (dashed green line) was used as a threshold to define the onset and offset of beta bursts (burst timings are shown in the green rectangular boxes). (c) Illustration of the fixed window approach, where 200 ms long data segments ending at fixed time intervals (0, 20, 40, 60, 80, or 100ms) prior to the onset of a burst are labelled as being predictive (Class 1) of subsequent burst onset. Non-predictive data segments (Class 0) terminated at least 150 ms prior to the onset of a burst. (d) Demonstration of the sliding window approach. A 200 ms long window with a stride length of 30 ms was passed along the beta filtered time series. 12 windowed segments with a shortened stride length of 5 ms (see main text) were labelled as being predictive of subsequent burst occurrence (Class 1). Windows that ended during the occurrence of a burst (Class Burst) were excluded from subsequent analysis, whilst the remaining data segments were labelled as being non-predictive (Class 0).

### Labelling of data for neural network classification

To train and test our network, we segmented beta band filtered STN timeseries using both sliding window and fixed window approaches. The fixed window approach allowed us to evaluate prediction performance at specific, consistent intervals leading up to the onset of bursts. In this approach, 200 ms long segments of the band filtered LFP, ending at designated intervals (0, 20, 40, 60, 80 or 100 ms) prior to the onset of each burst, were labelled as being predictive (**Class 1)** as per Figure 1c. Additionally, data terminating at least 150 ms before burst onset (excluding periods that coincided with burst timing) were divided into 200 ms long segments - with a stride of 50 ms - and categorized as non-predictive **(Class 0)**. During training, the number of Class 0 data segments was selected to match the number of Class 1 data segments. All segments were however classified during the validation and test phases.

The sliding window approach, using a 200 ms long window with stride lengths of 20, 25, or 30 ms, was designed to mimic real-time burst prediction for controlling the timing of stimulation delivery (Figure 1d shows an example with a stride length of 30 ms). If the end of a windowed segment overlapped with the occurrence of a burst, that segment was excluded from subsequent analysis (see **green** segments labelled **Class Burst** in Error! Reference source not found.**1d**). For training the network to learn burst predictive features, the final three data segments occurring prior to each burst were subsampled with a smaller stride length of 5 ms, yielding twelve 200 ms long data segments that terminated within 30-90 ms of burst onset (referred to as **Class 1** in **Figure 1d**). This smaller stride length served to expand the size of the training dataset and to allow the network to be sensitive to data features that exhibit subtly variable timing (smoothness) in relation to burst onset. The remaining 200 ms long data segments were associated with the absence of a subsequent burst and were labelled as being non-predictive (referred to as **Class 0** in **Figure 1d**).

The choice of sliding window length and stride length was made empirically, based on performance metrics from the validation datasets (see below section titled **Beta burst prediction network**). After testing three different window lengths (200, 250 and 300 ms) and seven stride lengths (10, 20, 25, 30, 35, 40, and 50 ms), we found that a 200 ms window with stride lengths between 20-30 ms resulted in optimal performance.

### Beta burst prediction network

The prediction network, illustrated in Figure 2, was constructed using a CNN architecture implemented in TensorFlow, using the Keras API (https://www.tensorflow.org and https://keras.io/api/). The LFP signal was passed through a sequence of 1D convolution, followed by a rectified linear unit (ReLU) activation function and 1D max pooling. This process was repeated three times to capture deep temporal features. The output from the final max pooling layer was flattened and connected to a dense layer, before a sigmoid function was used to perform binary classification (yielding an output of 0 or 1, corresponding to each of the classes). A kernel size of 5 was used for the 1D convolution layers, whilst a pooling size of 2 was used for the 1D max pooling layers to half the temporal dimensionality. The dense layer employed a dropout rate of 50%. The network was trained to minimize the categorical cross-entropy loss function, using the Adam optimizer with a learning rate of 0.0001^28^.

**Figure 2.**
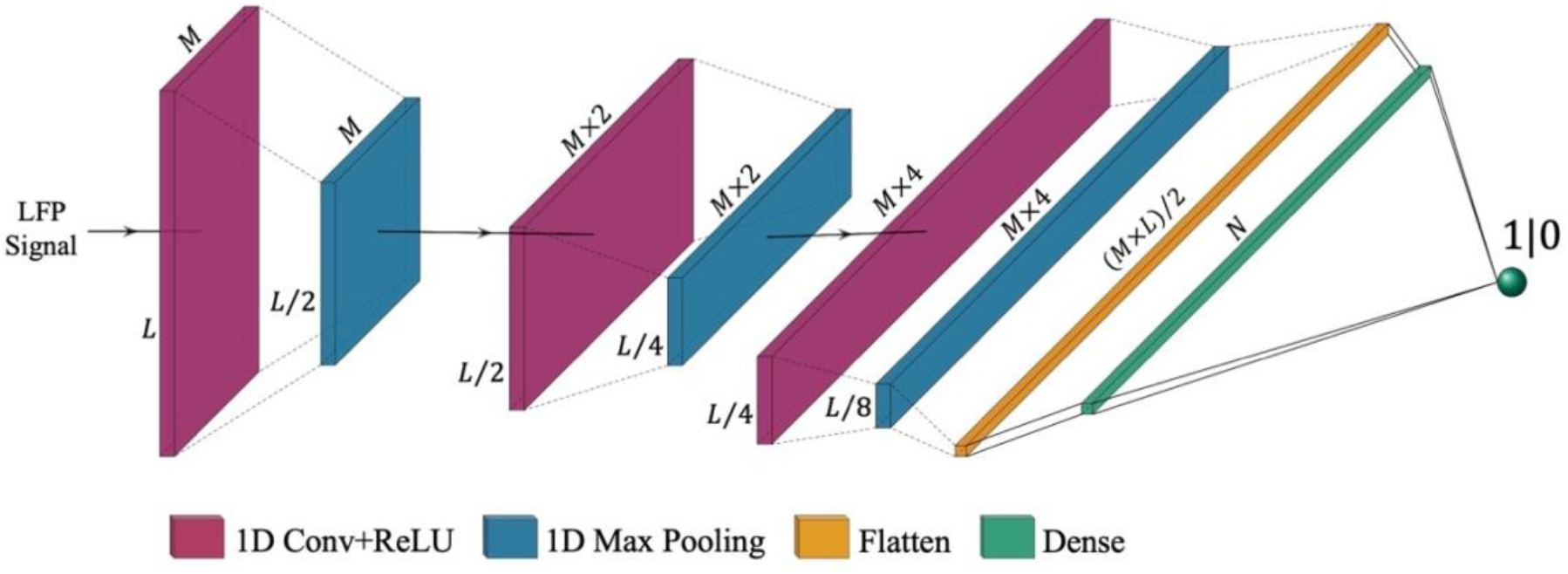
Architecture of the burst prediction network. The input beta filtered LFP is passed through a sequence of Conv1D, ReLU, and Max pooling (see main text). This process is repeated thrice, before the resulting outputs are flattened and passed to a dense layer. A single unit is then used for binary classification using a sigmoid activation function. In this analysis, data were segmented into 200 ms long epochs with a sampling rate of 600Hz, meaning that *L* = 120. *M* represents the number of filters in the convolutional layers and was selected to be 128. *N* indicates the number of neurons in the dense layer, *N=512*.

LFP data from each patient were split into training, validation, and test datasets. For patients with two recording sessions, one session was used exclusively for training, whilst the other was split evenly between validation and test datasets (50%-50%). For patients with only one recording session, the data were partitioned with 70% allocated for training, and 15% for each of validation and testing. The training data were used to fit multiple candidate models, while the validation set was utilized for selecting the best model architecture and optimizing hyperparameters (e.g., learning rate and the threshold of the final sigmoid output for binary classification). Final model performance was assessed and reported using the test dataset.

### Prediction performance metrics

Based on the comparison of test data labels and the model prediction, we calculated the following classification metrics:

- **True positives** (TP) – this is the number of burst predictive data segments (Class 1) that were correctly identified by the model.
- **True negatives** (TN) – this is the number of non-predictive data segments (Class 0) that were correctly identified by the model.
- **False positives** (FP) – this refers to non-predictive data segments (Class 0) where the model incorrectly predicted subsequent burst occurrence.
- **False negatives** (FN) – this refers to predictive data segments (Class 1), where the model failed to predict subsequent burst occurrence.
- **False positives per minute** (FP/min) – this quantifies the rate of false positive occurrence.
- **Accuracy** (ACC) – this indicates the proportion of correct predictions and is defined as: ACC = (TP+TN)/(TP+TN+FP+FN).
- **Precision** (PRC) – tells us the proportion of correct positive predictions and is defined as: PRC = TP/(TP+FP).
- **Sensitivity** (SEN) – also known as recall, this illustrates how well the model predicts the occurrence of bursts and is defined as: SEN=TP/(TP+FN).
- **Area under the receiver operating characteristic curve** (AUC-ROC) – this is a performance metric that considers the trade-off between sensitivity and the false positive rate at various thresholds. It provides a measure of the classifiers ability to correctly distinguish between classes and was used for the fixed window approach where class membership was balanced.
- **Area under the precision-recall curve** (AUC-PR) – this is a performance metric that considers the trade-off between precision and sensitivity at various thresholds and is therefore not influenced by a disproportionately high occurrence of true negative predictions ^29,30^. The AUC-PR was used instead of the AUC-ROC for the sliding window approach, owing to its increased effectiveness for classification evaluation on imbalanced datasets.
- **Prediction time prior to burst occurrence** (PT-PBO) – this metric was computed for the sliding window approach as the mean time interval between the end of the first true positive prediction (within 90 ms of burst onset) and the start of a burst.

## Results

### STN beta bursts can be predicted in advance of their onset

For the fixed window approach, we were able to achieve high burst prediction performance up to 60 ms before burst onset (mean values at -60 ms of: ACC = 0.78, SEN = 0.79, SPC = 0.78, AUC-PR = 0.73, AUC-ROC = 0.87). **Supplementary Tables 1-5** show burst prediction performance metrics for each patient, for each of the six different predictive window termination timepoints relative to burst onset (0, 20, 40, 60, 80 and 100 ms). This information is summarized in **Figure 3**, which shows mean estimates of ACC, SEN, SPC, AUC-PR, and AUC-ROC across patients for the six different predictive window termination timepoints.

**Figure 3.**
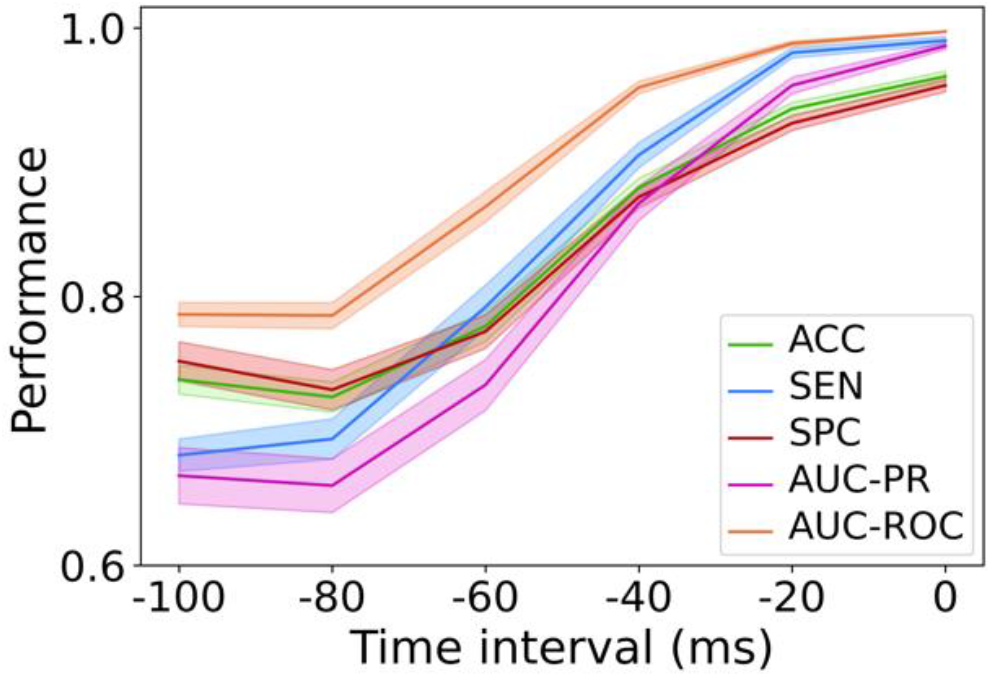
Burst prediction performance metrics for the fixed window approach. Results for each of the six different predictive window termination timepoints relative to burst onset are averaged across patients (ACC = accuracy; SEN = sensitivity; SPC = specificity; AUC-PR = area under the precision-recall curve; AUC-ROC = area under the ROC curve). Solid lines represent the mean, with the shaded regions indicating the standard error of the mean. Prediction performance metrics remain high as early as 60 ms prior to burst onset (see main Results for further discussion).

Individual patient prediction metrics for the sliding window approach, with a stride length of 30 ms, are shown in **Table 2**. Our model achieved a high performance for burst prediction, with a low false positive rate (mean sensitivity = 79%; mean precision = 88.8%; mean AUC-PR = 0.77; mean FP/min = 8.1). Importantly, the mean prediction time (PT-PBO) was 49.5 ms, which corresponds closely to the result obtained using the fixed window approach. The results of prediction performance using different stride lengths of 20 and 25 ms are shown in **Supplementary Tables 6-7**. Using these shorter stride lengths, it can be seen that prediction performance can be improved at the cost of a shorter prediction time (PT-PBO).

**Table 2.** Individual patient burst prediction performance metrics for the sliding window approach.

| Subject | Hemis<br>phere | TP | FP | FN | TN | SEN (%) | PRC (%) | AUC-PR | FP/<br>min | PT-PBO<br>(ms) |
| --- | --- | --- | --- | --- | --- | --- | --- | --- | --- | --- |
| 1 | R* | - | - | - | - | - | - | - | - | - |
|  | L | 111 | 14 | 34 | 2121 | 76.5 | 88.8 | 0.72 | 8.7 | 47.5 |
| 2 | R | 133 | 8 | 20 | 1842 | 86.9 | 94.3 | 0.81 | 4.9 | 48.0 |
|  | L | 121 | 13 | 24 | 1696 | 83.4 | 90.3 | 0.84 | 8.0 | 55.3 |
| 3 | R | 41 | 5 | 6 | 491 | 87.2 | 89.1 | 0.80 | 10.4 | 46.8 |
|  | L | 42 | 4 | 9 | 436 | 82.3 | 91.3 | 0.74 | 8.3 | 42.8 |
| 4 | R | 92 | 14 | 26 | 2171 | 78.0 | 86.8 | 0.78 | 8.9 | 55.1 |
|  | L | 101 | 16 | 41 | 2020 | 71.1 | 86.3 | 0.80 | 10.1 | 52.0 |
| 5 | R | 105 | 19 | 37 | 2757 | 73.9 | 84.6 | 0.73 | 9.5 | 49.1 |
|  | L | 101 | 10 | 38 | 1731 | 72.6 | 91.0 | 0.83 | 6.1 | 46.0 |
| 6 | R | 118 | 19 | 38 | 1676 | 75.6 | 86.1 | 0.81 | 12.0 | 52.6 |
|  | L | 110 | 22 | 33 | 2041 | 76.9 | 83.3 | 0.76 | 13.9 | 51.5 |
| 7 | R | 30 | 0 | 17 | 573 | 63.8 | 100 | 0.81 | 0 | 43.0 |
|  | L | 30 | 4 | 11 | 517 | 73.1 | 88.2 | 0.75 | 8.4 | 44.0 |
| 8 | R | 23 | 6 | 11 | 624 | 67.6 | 79.3 | 0.61 | 12.5 | 41.7 |
|  | L | 25 | 1 | 10 | 659 | 71.4 | 96.1 | 0.74 | 2.0 | 46.8 |
| 9 | R | 38 | 3 | 7 | 614 | 84.4 | 92.6 | 0.86 | 5.9 | 53.6 |
|  | L | 31 | 6 | 9 | 624 | 77.5 | 83.7 | 0.74 | 11.9 | 45.4 |
| 10 | R* | - | - | - | - | - | - | - | - | - |
|  | L | 47 | 4 | 5 | 489 | 90.3 | 92.1 | 0.77 | 8.3 | 44.6 |
| 11 | R | 67 | 13 | 11 | 2607 | 85.9 | 83.7 | 0.82 | 8.2 | 59.5 |
|  | L* | - | - | - | - | - | - | - | - | - |
| 12 | R | 44 | 4 | 2 | 832 | 95.6 | 91.6 | 0.82 | 6.3 | 50.4 |
|  | L | 47 | 1 | 6 | 772 | 88.6 | 97.9 | 0.84 | 1.5 | 54.2 |
| 13 | R | 32 | 4 | 13 | 577 | 71.1 | 88.8 | 0.76 | 8.5 | 46.8 |
|  | L | 34 | 2 | 15 | 530 | 69.3 | 94.4 | 0.73 | 4.2 | 45.5 |
| 14 | R | 25 | 3 | 8 | 723 | 75.7 | 89.2 | 0.78 | 5.94 | 64.8 |
|  | L | 33 | 4 | 4 | 593 | 89.1 | 89.1 | 0.81 | 7.9 | 55.4 |
| 15 | R | 26 | 4 | 7 | 489 | 78.7 | 86.6 | 0.72 | 8.4 | 47.3 |
|  | L | 26 | 7 | 3 | 597 | 89.6 | 78.7 | 0.77 | 14.7 | 55.3 |
| 16 | R | 151 | 22 | 38 | 2234 | 79.9 | 87.2 | 0.73 | 11.7 | 44.5 |
|  | L | 104 | 17 | 33 | 1730 | 75.9 | 85.9 | 0.76 | 10.6 | 47.3 |
| Sum | - | 1888 | 249 | 516 | 34766 | - | - | - | - | - |
| Mean±SEM | - | - | - | - | - | 79.0±2.1 | 88.8±1.3 | 0.77±0.0 | 8.1±1 | 49.5±1.5 |

\*These cases did not show a clear beta peak in the power spectrum and were therefore excluded from further analysis.

### Classification threshold controls the trade-off between sensitivity and false positive rate

An ideal burst prediction model should have a high sensitivity and a low false positive rate. The trade-off between these two performance metrics is determined by the sigmoid classification threshold of the output layer (see **Figure 2**), which binarises burst predictions. **Figure 4a and b** show the effect of varying the classification threshold on the SEN and FP rate for each patient’s burst prediction model, for the sliding window approach using a stride length of 30 ms. It is seen that lowering the threshold results in increased sensitivity at the cost of an increased FP rate. Analogous profiles for the sliding window approach using stride lengths of 20 and 25 ms are shown in **Supplementary Figure 1**.

**Figure 4.**
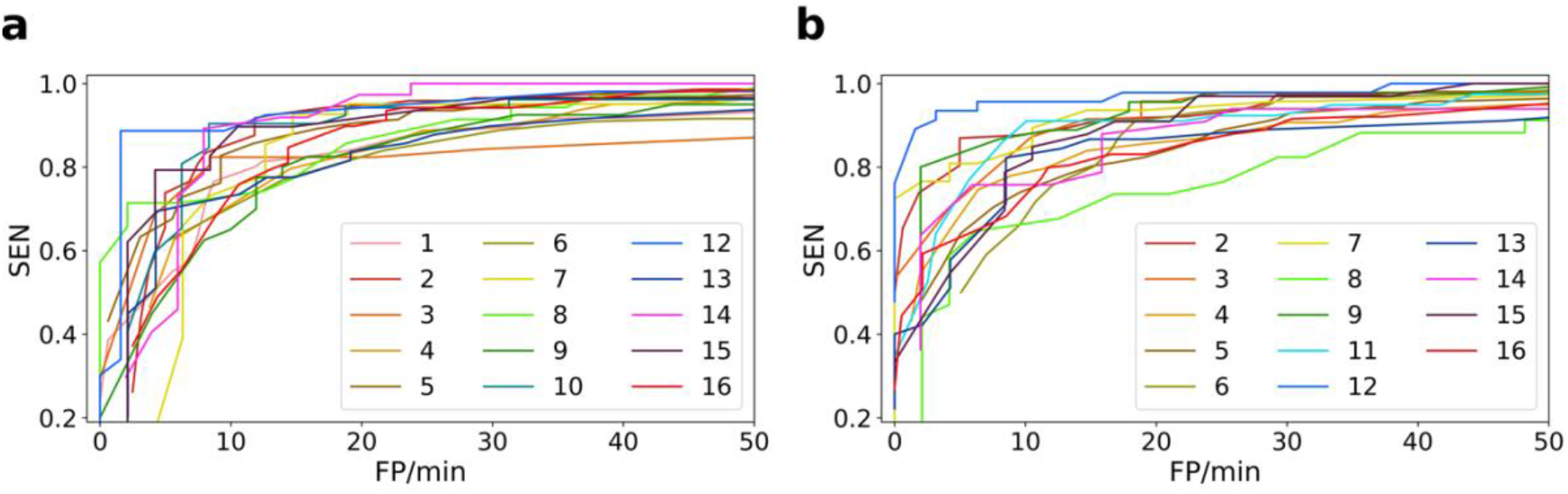
Classification threshold and the trade-off between sensitivity and false positive rate. Sensitivity (SEN) is plotted against the false positive rate (FP/min), for each patient’s prediction model, separately for the left (a) and right (b) hemispheric STN channels. Each coloured line represents a different patient. Lowering the classification threshold increases SEN, at the cost of an increased FP/min. Note that the left hemisphere of subject 11 and the right hemisphere of subjects 1 and 10 did not show a clear beta peak in the power spectrum and were therefore excluded from further analysis.

### Beta amplitude modulations predict subsequent burst occurrence

Our results reveal that beta burst occurrence may be accurately predicted as early as 60 ms before burst onset. But is there a consistent data feature allowing for this prediction which could be used in aDBS applications? To address this question, we examined mean beta amplitude envelopes for test data segments leading to TP, FP, and FN predictions. This procedure was performed separately for the right and left hemispheres, for each patient in the sliding window approach. In the case of TP and FN predictions, we selected the latest predictive data segment which terminated within 30 ms of burst onset (see **Figure 1**).

Figure 5. and **Figure 6** show the individual patient results of this analysis, separately for the right and left hemispheres. Crucially, the plots reveal that for TP predictions there is a consistent dip (fall followed by a subsequent rise) in the beta amplitude across patients, which occurs approximately 80-100 ms prior to beta burst onset. Note that in these figures the predictive window ends within 30 ms of burst onset. As expected, the beta amplitude profile of FP predictions closely matched that of TP predictions. FN predictions in contrast exhibited less pronounced beta amplitude dips. Taken together, these results highlight that a dip in the beta amplitude, occurring between 80-100 ms prior to burst onset, can be a reliable predictive biomarker of burst occurrence (see **Discussion** for further comments regarding this phenomenon).

### Validation of findings using surrogate data

We next sought to validate our findings, by generating surrogate data that preserved beta burst characteristics of the original data whilst destroying pathophysiological signal properties occurring within non-bursting time periods. We expected to see that this manipulation would lead to significantly impaired burst prediction performance metrics when training and testing our burst prediction network on surrogate data.

**Figure 5.**
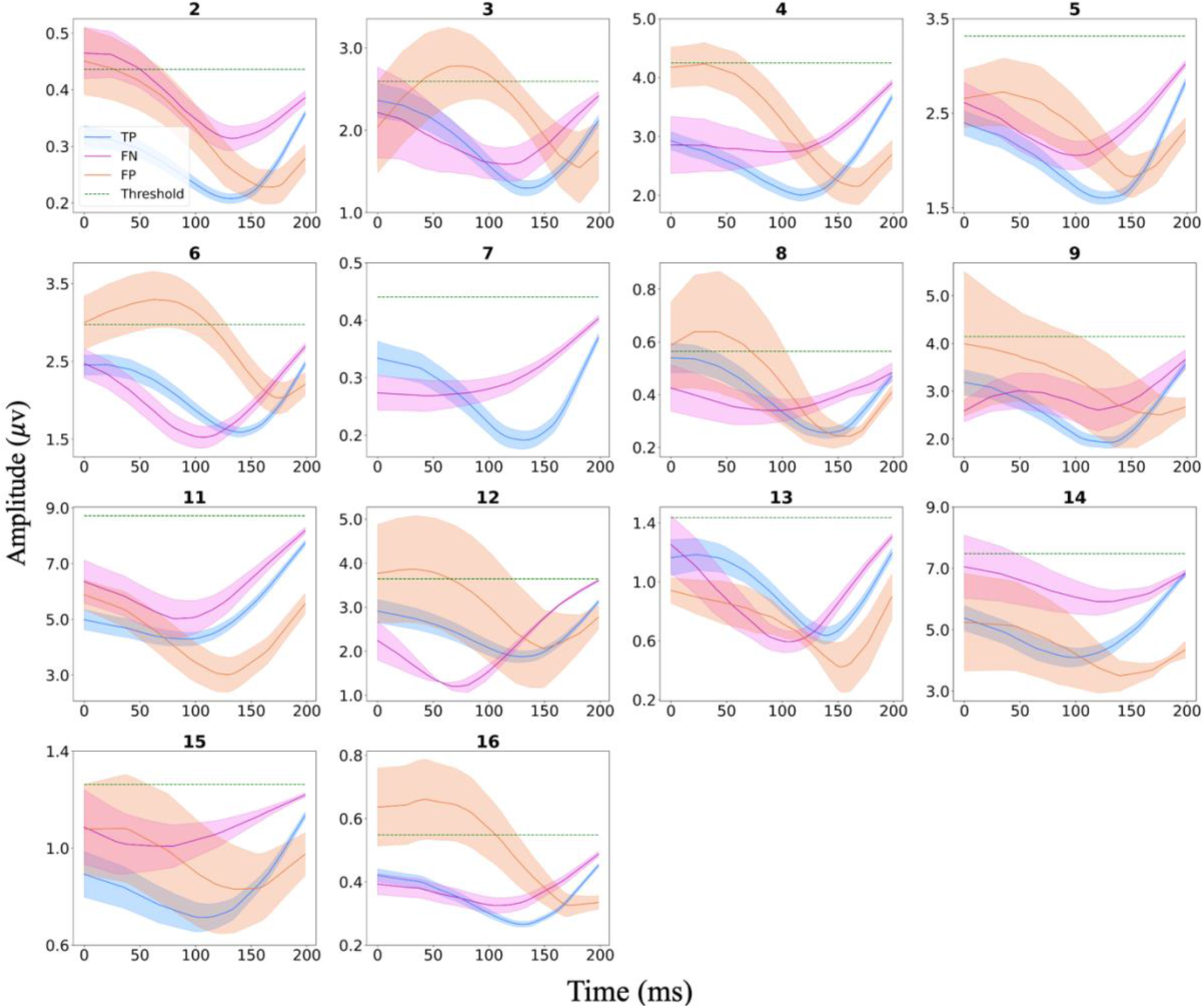
Individual patient beta envelope amplitudes for (200ms long) data segments leading to TP, TN, and FP predictions with the sliding window approach for the right hemispheric STN LFP. Solid lines indicate the mean, whilst shaded areas represent the standard error of the mean. For each patient the amplitude threshold (75^th^ percentile) for defining burst occurrence is also indicated by the green dashed line. For patient 7 there were no FP predictions. The right hemisphere of subjects 1 and 10 did not show a clear beta peak in the power spectrum and were therefore excluded from further analysis.

**Figure 6.**
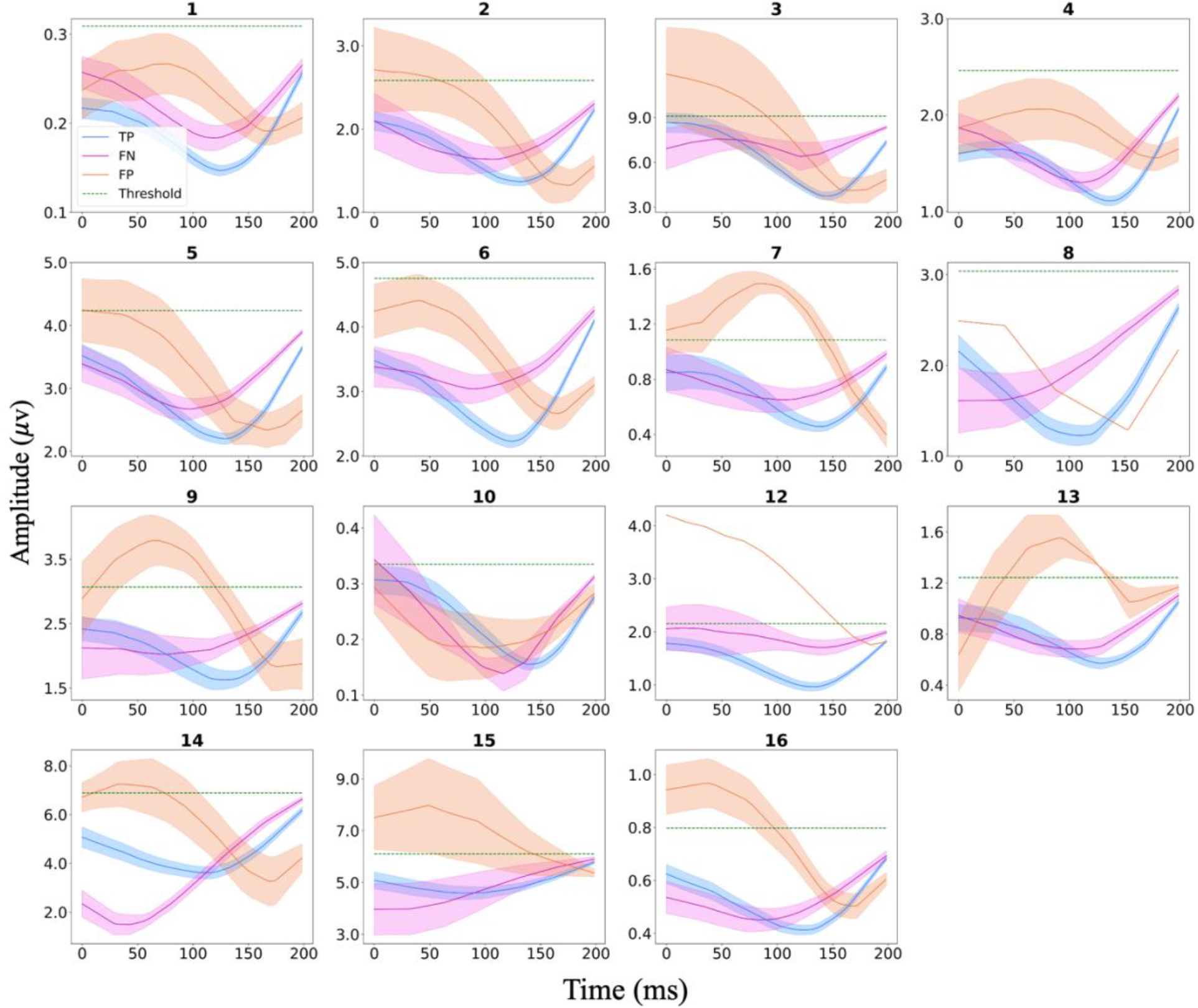
Individual patient beta envelope amplitudes for (200ms long) data segments leading to TP, TN, and FP predictions with the sliding window approach for the left hemisphere. Solid lines indicate the mean, whilst shaded areas represent the standard error of the mean. For each patient the amplitude threshold (75^th^ percentile) for defining burst occurrence is also indicated in the green dashed line. For patients 8 and 12, there was only one FP prediction. The left hemisphere of subject 11 did not show a clear beta peak in the power spectrum and was therefore excluded from further analysis.

The original signal from each patient and hemisphere was bandpass filtered (using a zero-phase 6^th^ order Butterworth filter with pass band of ±3 Hz around the beta peak frequency) to define periods of bursting activity. A surrogate signal was then created by preserving the original data from periods of bursting activity and randomly shuffling (in time) the data belonging to non-bursting time periods. The surrogate timeseries were then processed with the same causal filter used for the corresponding original dataset. A sliding window approach, with a stride length of 30 ms was employed to train and test the prediction network as before. **Supplementary Table 8** shows the performance of the prediction model using surrogate data. As anticipated, the model achieved poor performance compared to the original data, indicating that the network relies on the physiological signatures present in the original data to predict bursts, and that the narrow-band filter has no effect on training the network.

## Discussion

Our study demonstrates that the timing of pathological beta bursts within the parkinsonian STN can be accurately predicted from preceding STN beta activity. By training patient specific deep learning models, we show that bursts can be predicted with high sensitivity, accuracy, and precision up to 60 ms prior to their onset. Importantly, our approach requires only short amounts of training data - averaging less than 3 minutes for each patient – and can be deployed in real-time. Furthermore, by studying the characteristics of predictive data segments, we show that a dip in the beta amplitude, which occurs between 80-100 ms prior to burst onset, is a predictive biomarker of burst occurrence. These results provide proof-of-principle for the feasibility of burst prediction and also provide insights into the neurophysiological mechanisms leading to burst initiation.

### Translational potential of burst prediction

Beta frequency oscillatory activity within the parkinsonian cortico-basal ganglia circuit has proven to be a robust biomarker of motoric impairments – particularly bradykinesia and rigidity^4,7,33,34^. Moreover, aDBS approaches which involve the delivery of STN stimulation only after the detection of beta bursts are more effective in terms of achieving motor benefit and limiting stimulation related side effects than conventional continuous DBS^15–17,35^. Although aDBS has been shown to be effective, a key drawback of beta amplitude triggered approaches is that they cannot prevent the onset of pathological beta bursts, but rather respond with stimulation delivery after some fixed delay following burst onset. Consequently, aDBS may be acting too late to optimally prevent the initiation and propagation of pathological oscillatory activity within the cortico-basal ganglia circuit (akin to closing the stable door after the horse has bolted). This drawback speaks to the utility of patient specific burst prediction approaches which allow for the earlier detection of bursts, and consequently the earlier delivery of stimulation to facilitate their termination.

Although our findings provide the first demonstration of proof-of-principle, further real-time clinical testing of our approach in externalized patients is warranted before moving towards the implementation of prediction-based stimulation algorithms in DBS devices. Interestingly, our approach also draws important parallels with prior work exploring seizure forecasting from intracranial recordings. Deep learning-based approaches including CNNs have proven to be successful in this regard and may allow for the more rapid delivery of seizure abortive treatments such as medication or therapeutic stimulation ^36–38^. Theoretically, our approach may also be leveraged to predict other oscillatory features within and across frequency bands, which have been shown to relate to distinct cognitive and motor symptoms of PD ^32,39–41^, such as dyskinesia.

### Insights into network mechanisms leading to burst generation

A key finding of our work is that the beta amplitude displays a stereotyped pattern of modulation prior to the onset of bursts. **Figures 5 and 6** reveal the precise nature of this modulation, with the beta amplitude initially falling, reaching a nadir at approximately 80-100 ms prior to burst onset, and then subsequently rising again towards burst initiation. But what are the neural mechanisms which might explain this phenomenon? Computational models relying upon stochastic (noisy) inputs to neuronal populations as a primary driver of bursting cannot easily recapitulate the pre-burst amplitude dip ^18,42,43^. Usually in such models, noisy inputs determine transitions of the system between stable (non-oscillatory) and unstable (oscillatory) states via a bifurcation, meaning that oscillatory amplitudes can only change in one direction (either increase or decrease). A more plausible explanation for the beta amplitude dip might relate to the communication through coherence hypothesis, which proposes that maximal oscillatory entrainment within a network requires optimal phase alignment of interacting oscillatory populations ^44–46^. Switching to an optimal phase alignment for burst initiation requires a coordinated phase reset of oscillatory populations within the cortico-basal ganglia circuit, which could account for the transient decrease in the beta amplitude observed in our work ^46^.

### Study limitations

Our findings should be interpreted in the context of the following limitations. Firstly, STN recordings were performed a few days after electrode implantation, whilst patients were at rest and off dopaminergic medication. This meant that clinical constraints, including patient fatigue, allowed for only relatively short duration recordings. Nevertheless, our ongoing related work highlights that similar neural network architectures can be used to accurately predict burst properties across different behavioural and medication states in patients with sensing enabled DBS devices that allow for prolonged recordings ^47^. Importantly in this regard, the availability of longer training data can allow further improvements in prediction performance metrics. Finally, we have predicted STN beta bursts from STN activity and have therefore not included nodes within the cortico-basal ganglia circuit – such as primary motor cortex (M1) - whose activities may facilitate the prediction of STN bursts ^18,31^. Although electrocorticographic recordings from M1 are currently performed only for research purposes during DBS procedures, there is growing evidence to suggest that these may provide complementary information about clinical states (e.g., dyskinesia), particularly when the fidelity of subcortical recordings is compromised by the presence of stimulation artefacts^32^.

### Summary

We have harnessed deep neural networks to show that the timing of pathological beta bursts within the STN can be predicted in patients with PD. An important biomarker for this prediction is a dip in the beta amplitude, which occurs consistently between 80-100 ms prior to burst onset and may be indicative of a phase reset of oscillatory cortico-basal ganglia neuronal populations. Crucially, our findings motivate the development of a new generation of predictive DBS algorithms which are capable of either preventing pathological bursts or terminating them earlier following their onset.

## Acknowledgements

We thank Rafal Bogacz (University of Oxford) for helpful discussions regarding this work. This work was supported by an MRC Clinician Scientist Fellowship (MR/W024810/1) held by AO. BA and AO acknowledge funding support from the Oxford University Hospitals Charity. The Wellcome Centre for Human Neuroimaging is supported by core funding from Wellcome (203147/Z/16/Z). H.T. is supported by the Medical Research Council UK (MC_UU_00003/2, MR/V00655X/1, MR/P012272/1), the National Institute for Health Research (NIHR) Oxford Biomedical Research Centre (BRC) and the Rosetrees Trust, UK.

## Data and Code Availability

Anonymized datasets used for the current study and computer code are available from the corresponding authors on request and will be deposited on the MRC Brain Network Dynamics Unit data-sharing platform (https://data.mrc.ox.ac.uk/data-set/). Exemplar code for training and evaluation is available at https://github.com/b-abdi/Burst-Prediction/tree/main.

