## Supplementary Materials for "Prediction of pathological subthalamic nucleus beta burst occurrence in Parkinson’s disease"

**Supplementary Table 1**. Individual patient burst prediction accuracy (%) for the fixed window approach. The results were obtained for six different predictive window termination timepoints relative to burst onset (0, 20, 60, 80 and 100 ms).

| Sub | Hemisphere | -100ms | -80ms | -60ms | -40ms | -20ms | 0ms |
|---|---|---|---|---|---|---|---|
| 1 | R* | - | - | - | - | - | - |
| | L | 72.7 | 75.2 | 73.1 | 86.3 | 94 | 97.1 |
| 2 | R | 79.1 | 76.9 | 76.4 | 89.3 | 96.2 | 97.7 |
| | L | 78.8 | 74.9 | 85.3 | 90.4 | 95.1 | 98.2 |
| 3 | R | 77.4 | 65.3 | 78.9 | 89.4 | 96 | 95.5 |
| | L | 70.9 | 66.2 | 78.1 | 87.4 | 94.7 | 96 |
| 4 | R | 73 | 71.1 | 76.8 | 87.3 | 96.1 | 97.8 |
| | L | 73.3 | 72.1 | 81.6 | 92.5 | 96.3 | 99.1 |
| 5 | R | 78.1 | 79.3 | 83.8 | 90.8 | 94.9 | 96.3 |
| | L | 78.8 | 79.3 | 85 | 91.5 | 96.1 | 97.5 |
| 6 | R | 70.9 | 75.4 | 81.5 | 88.8 | 96 | 98.2 |
| | L | 73.7 | 73.2 | 80.4 | 90.1 | 94.9 | 96.6 |
| 7 | R | 72.2 | 69.7 | 70.7 | 84.8 | 90.9 | 94.9 |
| | L | 78 | 74 | 70.5 | 90.8 | 94.8 | 95.4 |
| 8 | R | 75.4 | 64.5 | 80.2 | 89.8 | 94.2 | 96.6 |
| | L | 61.1 | 65 | 69.7 | 83.3 | 93.6 | 95.3 |
| 9 | R | 74.7 | 72.2 | 86.1 | 90.7 | 96.2 | 99.2 |
| | L | 70.3 | 67.3 | 79.1 | 89.7 | 93.5 | 96.6 |
| 10 | R* | - | - | - | - | - | - |
| | L | 84.3 | 81.8 | 81.1 | 88.7 | 95.6 | 97.5 |
| 11 | R | 72.7 | 83.4 | 84.3 | 91.9 | 93.2 | 96.4 |
| | L* | - | - | - | - | - | - |
| 12 | R | 80.4 | 80.7 | 90.9 | 93.3 | 96.5 | 98 |
| | L | 71.6 | 68.1 | 79.8 | 92.9 | 96.8 | 99.6 |
| 13 | R | 73.3 | 77.4 | 71.9 | 84.6 | 89.1 | 91.4 |
| | L | 67.4 | 73.6 | 71.9 | 90.4 | 93.8 | 94.9 |
| 14 | R | 72.5 | 66.3 | 71.3 | 84.2 | 91.3 | 94.3 |
| | L | 74.9 | 70.5 | 87.7 | 93.4 | 96.9 | 98.2 |
| 15 | R | 50 | 51.5 | 61.3 | 76.3 | 85.6 | 89.2 |
| | L | 77.9 | 80.7 | 82.1 | 84.3 | 88.2 | 92.5 |
| 16 | R | 72.4 | 71.9 | 70.3 | 83.7 | 92.3 | 96.3 |
| | L | 77 | 69.7 | 76.8 | 87.6 | 92.7 | 95.6 |
| **Mean** | **-** | **73.5** | **72.3** | **78.1** | **88.4** | **94.0** | **96.3** |

*These cases did not show a clear beta peak in the power spectrum and were therefore excluded from further analysis.

**Supplementary Table 2**. Individual patient burst prediction sensitivity (%) for the fixed window approach. The results were obtained for six different predictive window termination timepoints relative to burst onset (0, 20, 60, 80 and 100ms).

| Sub | Hemisphere | -100ms | -80ms | -60ms | -40ms | -20ms | 0ms |
|---|---|---|---|---|---|---|---|
| 1 | R* | - | - | - | - | - | - |
| | L | 59.4 | 57.3 | 60.8 | 89.5 | 97.2 | 99.3 |
| 2 | R | 66 | 64.7 | 69.9 | 92.8 | 100 | 100 |
| | L | 65.5 | 66.2 | 91.2 | 97.3 | 99.3 | 100 |
| 3 | R | 66.7 | 68.9 | 88.9 | 91.1 | 97.8 | 100 |
| | L | 81.1 | 90.6 | 94.3 | 98.1 | 98.1 | 100 |
| 4 | R | 58.8 | 63 | 82.4 | 94.1 | 98.3 | 99.2 |
| | L | 66.9 | 64.9 | 82.4 | 95.3 | 99.3 | 100 |
| 5 | R | 63.6 | 69.9 | 79.7 | 88.1 | 98.6 | 99.3 |
| | L | 73.6 | 69.4 | 81.2 | 93.1 | 98.6 | 98.6 |
| 6 | R | 72.9 | 71.7 | 85.5 | 91 | 97.6 | 100 |
| | L | 68.8 | 68.1 | 77.3 | 85.1 | 95 | 99.3 |
| 7 | R | 64.4 | 60 | 55.6 | 84.4 | 100 | 97.8 |
| | L | 61.9 | 59.5 | 71.4 | 92.9 | 100 | 100 |
| 8 | R | 73.7 | 78.9 | 84.2 | 86.8 | 92.1 | 94.7 |
| | L | 60 | 71.4 | 71.4 | 77.1 | 100 | 91.4 |
| 9 | R | 66.7 | 64.4 | 75.6 | 82.2 | 95.6 | 97.8 |
| | L | 68.9 | 77.8 | 77.8 | 86.7 | 97.8 | 100 |
| 10 | R* | - | - | - | - | - | - |
| | L | 66.1 | 72.9 | 71.2 | 86.4 | 98.3 | 98.3 |
| 11 | R | 46.7 | 42.7 | 88 | 98.7 | 100 | 100 |
| | L* | - | - | - | - | - | - |
| 12 | R | 74.5 | 72.3 | 93.6 | 97.9 | 100 | 100 |
| | L | 77.4 | 79.2 | 92.5 | 96.2 | 100 | 100 |
| 13 | R | 71.1 | 71.1 | 71.1 | 93.3 | 100 | 100 |
| | L | 72.9 | 81.2 | 77.1 | 91.7 | 100 | 100 |
| 14 | R | 63.3 | 70 | 66.7 | 93.3 | 100 | 100 |
| | L | 70.3 | 70.3 | 89.2 | 94.6 | 100 | 100 |
| 15 | R | 74.2 | 64.5 | 77.4 | 83.9 | 93.5 | 100 |
| | L | 79.3 | 82.8 | 89.7 | 89.7 | 93.1 | 100 |
| 16 | R | 71.1 | 67.5 | 75.8 | 84 | 96.4 | 99 |
| | L | 65.8 | 67.1 | 80.1 | 88.4 | 97.9 | 97.3 |
| Mean | - | 68.0 | 69.3 | 79.4 | 90.5 | 98.1 | 99.0 |

*These cases did not show a clear beta peak in the power spectrum and were therefore excluded from further analysis.

**Supplementary Table 3**. Individual patient burst prediction specificity (%) for the fixed window approach. The results were obtained for six different predictive window termination timepoints relative to burst onset (0, 20, 60, 80 and 100ms).

| Sub | Hemisphere | -100ms | -80ms | -60ms | -40ms | -20ms | 0ms |
|---|---|---|---|---|---|---|---|
| 1 | R* | - | - | - | - | - | - |
| | L | 75.3 | 78.7 | 75.4 | 85.7 | 93.4 | 96.6 |
| 2 | R | 82.7 | 80.3 | 78.1 | 88.3 | 95.1 | 97.1 |
| | L | 83 | 77.6 | 83.4 | 88.2 | 93.8 | 97.6 |
| 3 | R | 80.5 | 64.3 | 76 | 89 | 95.5 | 94.2 |
| | L | 65.3 | 53.1 | 69.4 | 81.6 | 92.9 | 93.9 |
| 4 | R | 75 | 72.3 | 76 | 86.3 | 95.8 | 97.7 |
| | L | 74.8 | 73.8 | 81.3 | 91.8 | 95.6 | 98.9 |
| 5 | R | 80 | 80.6 | 84.4 | 91.2 | 94.4 | 95.9 |
| | L | 80.4 | 82.2 | 86 | 91.1 | 95.3 | 97.2 |
| 6 | R | 70.2 | 76.8 | 80.1 | 88 | 95.4 | 97.6 |
| | L | 74.7 | 74.3 | 81.1 | 91.2 | 94.9 | 96.1 |
| 7 | R | 74.5 | 72.5 | 75.2 | 85 | 88.2 | 94.1 |
| | L | 83.2 | 78.6 | 70.2 | 90.1 | 93.1 | 93.9 |
| 8 | R | 75.7 | 62.4 | 79.6 | 90.2 | 94.5 | 96.9 |
| | L | 61.3 | 63.8 | 69.3 | 84.4 | 92.5 | 96 |
| 9 | R | 76.6 | 74 | 88.5 | 92.7 | 96.4 | 99.5 |
| | L | 70.6 | 65.1 | 79.4 | 90.4 | 92.7 | 95.9 |
| 10 | R* | - | - | - | - | - | - |
| | L | 95 | 87 | 87 | 90 | 94 | 97 |
| 11 | R | 74.4 | 86.1 | 84 | 91.4 | 92.8 | 96.2 |
| | L* | - | - | - | - | - | - |
| 12 | R | 81.4 | 82 | 90.5 | 92.5 | 95.9 | 97.6 |
| | L | 70.3 | 65.5 | 76.9 | 92.1 | 96.1 | 99.6 |
| 13 | R | 73.9 | 79 | 72.2 | 82.4 | 86.4 | 89.2 |
| | L | 65.4 | 70.8 | 70 | 90 | 91.5 | 93.1 |
| 14 | R | 73.4 | 65.9 | 71.8 | 83.3 | 90.5 | 93.8 |
| | L | 75.8 | 70.5 | 87.4 | 93.2 | 96.3 | 97.9 |
| 15 | R | 45.4 | 49.1 | 58.3 | 74.8 | 84 | 87.1 |
| | L | 77.7 | 80.5 | 81.3 | 83.7 | 87.6 | 91.6 |
| 16 | R | 72.8 | 73.2 | 68.6 | 83.7 | 91 | 95.5 |
| | L | 79.9 | 70.3 | 76 | 87.5 | 91.3 | 95.2 |
| **Mean** | **-** | **74.8** | **72.8** | **77.8** | **87.9** | **93.0** | **95.6** |

*These cases did not show a clear beta peak in the power spectrum and were therefore excluded from further analysis.

**Supplementary Table 4**. Individual patient burst prediction AUC-PR for the fixed window approach. The results were obtained for six different predictive window termination timepoints relative to burst onset (0, 20, 60, 80 and 100ms).

| Sub | Hemisphere | -100ms | -80ms | -60ms | -40ms | -20ms | 0ms |
|---|---|---|---|---|---|---|---|
| 1 | R* | - | - | - | - | - | - |
| | L | 0.59 | 0.58 | 0.58 | 0.83 | 0.96 | 0.99 |
| 2 | R | 0.7 | 0.68 | 0.72 | 0.91 | 0.99 | 1 |
| | L | 0.73 | 0.72 | 0.87 | 0.96 | 0.99 | 1 |
| 3 | R | 0.75 | 0.72 | 0.84 | 0.92 | 0.98 | 1 |
| | L | 0.87 | 0.86 | 0.88 | 0.95 | 0.99 | 1 |
| 4 | R | 0.52 | 0.53 | 0.64 | 0.81 | 0.94 | 0.98 |
| | L | 0.66 | 0.65 | 0.79 | 0.93 | 0.98 | 1 |
| 5 | R | 0.6 | 0.6 | 0.71 | 0.82 | 0.93 | 0.98 |
| | L | 0.76 | 0.75 | 0.84 | 0.92 | 0.98 | 0.99 |
| 6 | R | 0.75 | 0.75 | 0.86 | 0.94 | 0.98 | 1 |
| | L | 0.62 | 0.6 | 0.71 | 0.84 | 0.93 | 0.98 |
| 7 | R | 0.68 | 0.65 | 0.59 | 0.8 | 0.96 | 0.97 |
| | L | 0.68 | 0.66 | 0.67 | 0.92 | 0.98 | 1 |
| 8 | R | 0.67 | 0.67 | 0.78 | 0.86 | 0.94 | 0.99 |
| | L | 0.5 | 0.48 | 0.54 | 0.69 | 0.91 | 0.95 |
| 9 | R | 0.69 | 0.65 | 0.76 | 0.88 | 0.96 | 1 |
| | L | 0.63 | 0.66 | 0.73 | 0.85 | 0.94 | 1 |
| 10 | R* | - | - | - | - | - | 1 |
| | L | 0.87 | 0.85 | 0.87 | 0.93 | 0.99 | 1 |
| 11 | R | 0.28 | 0.3 | 0.64 | 0.85 | 0.9 | 0.96 |
| | L* | - | - | - | - | - | 1 |
| 12 | R | 0.7 | 0.72 | 0.89 | 0.97 | 1 | 1 |
| | L | 0.71 | 0.72 | 0.8 | 0.95 | 0.99 | 1 |
| 13 | R | 0.68 | 0.71 | 0.66 | 0.85 | 0.98 | 0.99 |
| | L | 0.74 | 0.76 | 0.75 | 0.94 | 0.95 | 0.99 |
| 14 | R | 0.45 | 0.49 | 0.47 | 0.75 | 0.94 | 0.97 |
| | L | 0.65 | 0.63 | 0.83 | 0.9 | 0.97 | 0.97 |
| 15 | R | 0.52 | 0.48 | 0.57 | 0.67 | 0.83 | 0.95 |
| | L | 0.6 | 0.63 | 0.69 | 0.79 | 0.9 | 0.96 |
| 16 | R | 0.71 | 0.69 | 0.71 | 0.83 | 0.96 | 0.99 |
| | L | 0.69 | 0.65 | 0.74 | 0.85 | 0.96 | 0.98 |
| **Mean** | - | **0.66** | **0.65** | **0.73** | **0.87** | **0.96** | **0.99** |

*These cases did not show a clear beta peak in the power spectrum and were therefore excluded from further analysis.

**Supplementary Table 5**. Individual patient burst prediction AUC-ROC for the fixed window approach. The results were obtained for six different predictive window termination timepoints relative to burst onset (0, 20, 60, 80 and 100ms).

| Sub | Hemisphere | -100ms | -80ms | -60ms | -40ms | -20ms | 0ms |
|---|---|---|---|---|---|---|---|
| 1 | R* | - | - | - | - | - | - |
| | L | 0.73 | 0.74 | 0.75 | 0.95 | 0.99 | 1 |
| 2 | R | 0.77 | 0.76 | 0.82 | 0.97 | 1 | 1 |
| | L | 0.8 | 0.8 | 0.94 | 0.98 | 1 | 1 |
| 3 | R | 0.81 | 0.78 | 0.91 | 0.97 | 0.99 | 1 |
| | L | 0.87 | 0.87 | 0.93 | 0.97 | 0.99 | 1 |
| 4 | R | 0.72 | 0.75 | 0.88 | 0.97 | 0.99 | 1 |
| | L | 0.78 | 0.78 | 0.91 | 0.98 | 1 | 1 |
| 5 | R | 0.78 | 0.81 | 0.9 | 0.97 | 0.99 | 1 |
| | L | 0.83 | 0.84 | 0.92 | 0.97 | 0.99 | 1 |
| 6 | R | 0.79 | 0.81 | 0.92 | 0.97 | 0.99 | 1 |
| | L | 0.79 | 0.77 | 0.87 | 0.96 | 0.99 | 1 |
| 7 | R | 0.78 | 0.71 | 0.69 | 0.91 | 0.99 | 0.99 |
| | L | 0.74 | 0.7 | 0.81 | 0.97 | 0.99 | 1 |
| 8 | R | 0.81 | 0.82 | 0.92 | 0.94 | 0.96 | 1 |
| | L | 0.7 | 0.71 | 0.8 | 0.91 | 0.98 | 0.99 |
| 9 | R | 0.79 | 0.74 | 0.89 | 0.95 | 0.97 | 1 |
| | L | 0.76 | 0.8 | 0.87 | 0.95 | 0.99 | 1 |
| 10 | R* | - | - | - | - | - | - |
| | L | 0.87 | 0.85 | 0.88 | 0.94 | 0.99 | 1 |
| 11 | R | 0.65 | 0.68 | 0.95 | 0.99 | 0.99 | 1 |
| | L* | - | - | - | - | - | - |
| 12 | R | 0.84 | 0.84 | 0.97 | 0.99 | 1 | 1 |
| | L | 0.82 | 0.82 | 0.92 | 0.99 | 1 | 1 |
| 13 | R | 0.81 | 0.83 | 0.81 | 0.95 | 0.99 | 1 |
| | L | 0.8 | 0.84 | 0.84 | 0.97 | 0.99 | 1 |
| 14 | R | 0.74 | 0.79 | 0.8 | 0.96 | 0.99 | 1 |
| | L | 0.83 | 0.79 | 0.94 | 0.96 | 1 | 1 |
| 15 | R | 0.68 | 0.66 | 0.79 | 0.88 | 0.95 | 0.99 |
| | L | 0.84 | 0.89 | 0.94 | 0.96 | 0.98 | 0.99 |
| 16 | R | 0.8 | 0.78 | 0.8 | 0.91 | 0.98 | 1 |
| | L | 0.77 | 0.75 | 0.87 | 0.94 | 0.99 | 1 |
| **Mean** | **-** | **0.78** | **0.78** | **0.87** | **0.96** | **0.99** | **1** |

*These cases did not show a clear beta peak in the power spectrum and were therefore excluded from further analysis.

**Supplementary Table 6.** Individual patient burst prediction performance metrics for the sliding window approach with a stride length of 20 ms.

| Subject | Hemisphere | TP | FP | FN | TN | SEN (%) | PRC (%) | AUC-PR | FP/min | PT-PBO |
|---|---|---|---|---|---|---|---|---|---|---|
| 1 | R* | - | - | - | - | - | - | - | - | - |
| | L | 118 | 11 | 34 | 3390 | 77.6 | 91.4 | 83.2 | 6.8 | 35.0 |
| 2 | R | 131 | 14 | 27 | 2981 | 82.9 | 90.3 | 85.8 | 8.7 | 39.0 |
| | L | 128 | 17 | 21 | 2751 | 85.9 | 88.2 | 90.5 | 10.5 | 41.4 |
| 3 | R | 42 | 2 | 6 | 809 | 87.5 | 95.4 | 90.7 | 4.1 | 38.5 |
| | L | 45 | 3 | 10 | 721 | 81.8 | 93.7 | 87.5 | 6.2 | 41.1 |
| 4 | R | 101 | 16 | 19 | 3435 | 84.1 | 86.3 | 82.2 | 10.2 | 38.8 |
| | L | 109 | 7 | 38 | 3234 | 74.1 | 93.9 | 88.5 | 4.4 | 36.8 |
| 5 | R | 118 | 12 | 27 | 4349 | 81.3 | 90.7 | 84.9 | 6.0 | 33.9 |
| | L | 123 | 14 | 22 | 2797 | 84.8 | 89.7 | 86.5 | 8.6 | 38.2 |
| 6 | R | 142 | 11 | 22 | 2741 | 86.5 | 92.8 | 90.3 | 6.9 | 37.7 |
| | L | 120 | 8 | 27 | 3286 | 81.6 | 93.7 | 86.9 | 5.0 | 37.1 |
| 7 | R | 41 | 0 | 6 | 930 | 87.2 | 100 | 87.8 | 0 | 36.5 |
| | L | 40 | 1 | 4 | 834 | 90.9 | 97.5 | 90.2 | 2.1 | 38.0 |
| 8 | R | 23 | 4 | 14 | 976 | 62.1 | 85.1 | 74.2 | 8.3 | 35.5 |
| | L | 27 | 5 | 8 | 1035 | 77.1 | 84.3 | 80.6 | 10.4 | 40.0 |
| 9 | R | 40 | 2 | 6 | 990 | 86.9 | 95.2 | 91.6 | 3.9 | 44.0 |
| | L | 40 | 1 | 4 | 834 | 90.9 | 97.5 | 90.2 | 2.0 | 38.0 |
| 10 | R* | - | - | - | - | - | - | - | - | - |
| | L | 51 | 2 | 3 | 797 | 94.4 | 96.2 | 90.2 | 4.1 | 39.2 |
| 11 | R | 65 | 12 | 14 | 4028 | 82.2 | 84.4 | 87.0 | 7.5 | 42.7 |
| | L* | - | - | - | - | - | - | - | - | - |
| 12 | R | 45 | 5 | 1 | 1320 | 97.8 | 90.0 | 89.7 | 7.9 | 43.1 |
| | L | 49 | 1 | 5 | 1233 | 90.7 | 98.0 | 92.4 | 1.5 | 47.7 |
| 13 | R | 41 | 5 | 5 | 925 | 89.1 | 89.1 | 83.4 | 10.6 | 38.0 |
| | L | 46 | 2 | 6 | 867 | 88.4 | 95.8 | 88.3 | 4.2 | 39.1 |
| 14 | R | 28 | 2 | 6 | 1128 | 82.3 | 93.3 | 84.7 | 3.9 | 40.0 |
| | L | 38 | 4 | 2 | 934 | 95.0 | 90.4 | 89.6 | 7.9 | 45.7 |
| 15 | R | 28 | 3 | 5 | 784 | 84.8 | 90.3 | 81.1 | 6.3 | 37.1 |
| | L | 24 | 4 | 5 | 945 | 82.7 | 85.7 | 78.3 | 8.4 | 38.3 |
| 16 | R | 177 | 12 | 21 | 3635 | 89.3 | 93.6 | 86.1 | 6.4 | 37.5 |
| | L | 124 | 11 | 27 | 2784 | 82.1 | 91.8 | 84.8 | 6.8 | 36.1 |
| **Sum** | **-** | **2104** | **191** | **395** | **55473** | - | - | - | - | - |
| **Mean** | **-** | **-** | **-** | **-** | **-** | **84.9** | **91.8** | **86.4** | **6.1** | **39.1** |

*These cases did not show a clear beta peak in the power spectrum and were therefore excluded from further analysis.

**Supplementary Table 7.** Individual patient burst prediction performance metrics for the sliding window approach with a stride length of 25 ms.

| Subject | Hemisphere | TP | FP | FN | TN | SEN (%) | PRC (%) | AUC-PR | FP/min | PT-PBO |
|---|---|---|---|---|---|---|---|---|---|---|
| 1 | R* | - | - | - | - | - | - | - | - | - |
| | L | 116 | 12 | 32 | 2618 | 78.3 | 90.6 | 80.6 | 7.4 | 42.0 |
| 2 | R | 139 | 14 | 19 | 2280 | 88.0 | 90.8 | 83.7 | 8.7 | 41.3 |
| | L | 126 | 13 | 22 | 2114 | 85.1 | 90.6 | 88.2 | 80.0 | 48.6 |
| 3 | R | 40 | 2 | 8 | 616 | 83.3 | 95.2 | 87.3 | 4.1 | 41.2 |
| | L | 40 | 3 | 13 | 552 | 75.4 | 93.0 | 84.0 | 6.2 | 42.7 |
| 4 | R | 97 | 8 | 21 | 2683 | 82.2 | 92.3 | 82.2 | 5.1 | 45.3 |
| | L | 114 | 18 | 29 | 2507 | 79.7 | 86.3 | 84.9 | 11.3 | 45.6 |
| 5 | R | 112 | 15 | 33 | 3393 | 77.2 | 88.1 | 80.0 | 7.5 | 46.2 |
| | L | 108 | 9 | 37 | 2150 | 74.4 | 92.3 | 86.5 | 5.5 | 43.2 |
| 6 | R | 135 | 16 | 24 | 2106 | 84.9 | 89.4 | 87.0 | 10.1 | 46.4 |
| | L | 113 | 8 | 32 | 2544 | 78.0 | 93.3 | 84.7 | 5.0 | 47.5 |
| 7 | R | 31 | 1 | 16 | 711 | 65.9 | 96.8 | 82.1 | 2.1 | 43.5 |
| | L | 39 | 2 | 3 | 646 | 92.8 | 95.1 | 82.6 | 4.2 | 37.8 |
| 8 | R | 27 | 6 | 8 | 766 | 77.1 | 81.8 | 67.2 | 12.5 | 38.8 |
| | L | 26 | 5 | 9 | 810 | 74.2 | 83.8 | 79.5 | 10.4 | 46.1 |
| 9 | R | 33 | 2 | 12 | 746 | 73.3 | 94.2 | 85.8 | 3.9 | 47.7 |
| | L | 39 | 3 | 5 | 761 | 88.6 | 92.8 | 85.1 | 5.9 | 47.4 |
| 10 | R* | - | - | - | - | - | - | - | - | - |
| | L | 44 | 7 | 8 | 610 | 84.6 | 86.2 | 77.4 | 14.5 | 39.2 |
| 11 | R | 65 | 11 | 14 | 3180 | 82.2 | 85.5 | 84.0 | 6.9 | 51.9 |
| | L* | - | - | - | - | - | - | - | - | - |
| 12 | R | 44 | 5 | 2 | 1028 | 95.6 | 89.8 | 87.4 | 7.9 | 48.3 |
| | L | 47 | 1 | 6 | 951 | 88.6 | 97.9 | 91.0 | 1.5 | 47.3 |
| 13 | R | 33 | 3 | 13 | 714 | 71.7 | 91.6 | 81.6 | 6.3 | 46.2 |
| | L | 38 | 4 | 13 | 660 | 74.5 | 90.4 | 82.9 | 8.5 | 41.4 |
| 14 | R | 27 | 5 | 7 | 880 | 79.4 | 84.3 | 80.2 | 9.9 | 48.1 |
| | L | 34 | 4 | 5 | 726 | 87.1 | 89.4 | 85.7 | 7.9 | 53.6 |
| 15 | R | 27 | 0 | 6 | 609 | 81.8 | 100 | 77.7 | 0 | 32.4 |
| | L | 25 | 3 | 4 | 737 | 86.2 | 89.2 | 80.3 | 6.3 | 48.0 |
| 16 | R | 146 | 8 | 48 | 2793 | 75.2 | 94.8 | 79.2 | 4.2 | 36.8 |
| | L | 97 | 7 | 52 | 2138 | 65.1 | 93.2 | 80.3 | 4.3 | 41.7 |
| **Sum** | **-** | **1962** | **195** | **501** | **43029** | - | - | - | - | - |
| **Mean** | **-** | **-** | **-** | **-** | **-** | **80.3** | **91.0** | **82.7** | **9.2** | **44.3** |

*These cases did not show a clear beta peak in the power spectrum and were therefore excluded from further analysis.

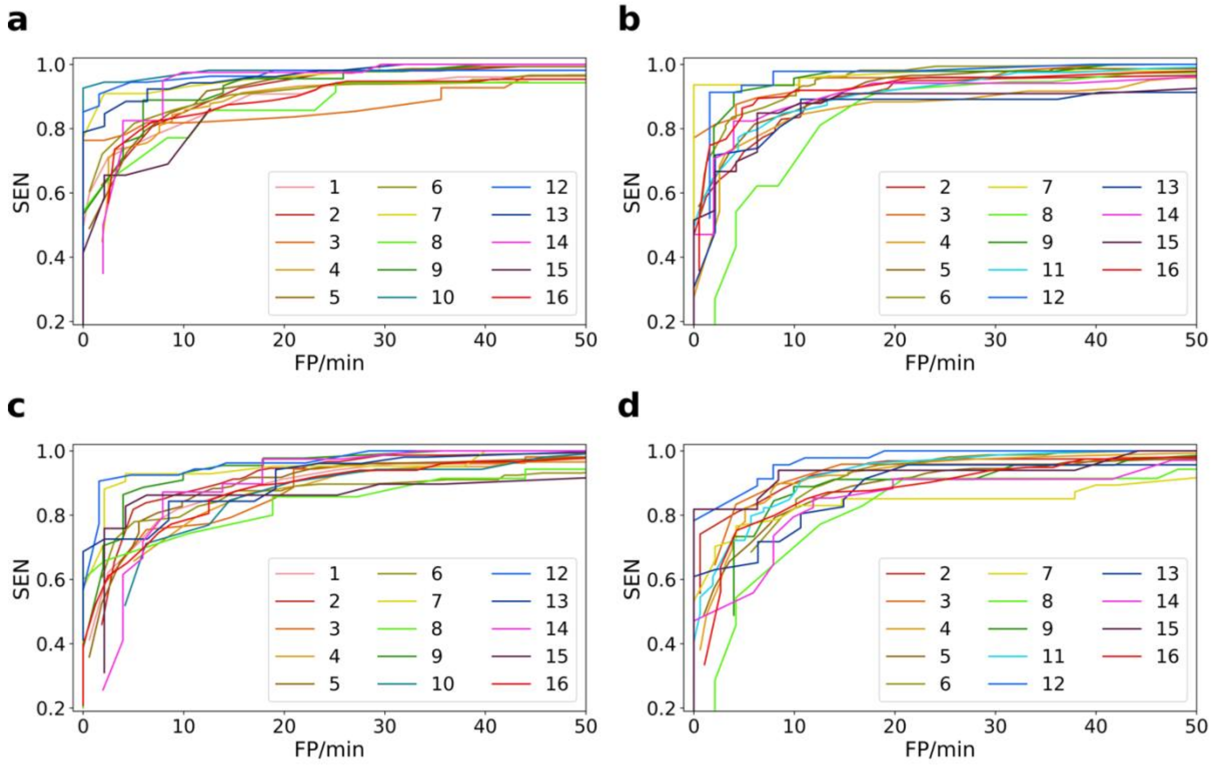


**Supplementary Figure 1.** The trade-off between sensitivity and false positive rate (FP/min) obtained using the sliding window approach is shown for: (a) left hemispheric STN channels with a stride length of 20 ms, (b) right hemispheric STN channels with a stride length of 20 ms, (c) left hemispheric STN channels with a stride length of 25 ms, and (d) right hemispheric STN channels with a stride length of 25 ms. Each coloured line represents a different patient. Note that the left hemisphere of subject 11 and the right hemisphere of subjects 1 and 10 did not show a clear beta peak in the power spectrum and were therefore excluded from further analysis.

**Supplementary Table 8.** The performance of model obtained using the surrogate data.

| Subject | Hemisphere | TP | FP | FN | TN | SEN (%) | PRC (%) | AUC-PR | FP/min | PT-PBO |
|---|---|---|---|---|---|---|---|---|---|---|
| 1 | R* | - | - | - | - | - | - | - | - | - |
| | L | 37 | 10 | 108 | 2125 | 25.5 | 78.7 | 0.46 | 6.2 | 54 |
| 2 | R | 41 | 18 | 112 | 1832 | 26.7 | 69.4 | 0.45 | 11.1 | 46 |
| | L | 81 | 21 | 64 | 1688 | 55.8 | 79.4 | 0.64 | 13.0 | 58 |
| 3 | R | 15 | 5 | 32 | 491 | 31.9 | 75.0 | 0.48 | 10.4 | 46 |
| | L | 39 | 5 | 12 | 435 | 76.4 | 88.6 | 0.78 | 10.4 | 51 |
| 4 | R | 81 | 17 | 37 | 2168 | 68.6 | 82.6 | 0.72 | 10.8 | 60.7 |
| | L | 33 | 15 | 109 | 2021 | 23.2 | 68.7 | 0.48 | 9.4 | 50 |
| 5 | R | 88 | 24 | 54 | 2752 | 61.9 | 78.5 | 0.66 | 12.1 | 52 |
| | L | 64 | 17 | 75 | 1724 | 46 | 79.0 | 0.53 | 10.4 | 54.8 |
| 6 | R | 58 | 17 | 98 | 1678 | 37.1 | 77.3 | 0.51 | 10.7 | 48.1 |
| | L | 75 | 16 | 68 | 2047 | 52.4 | 82.4 | 0.63 | 10.1 | 55.9 |
| 7 | R | 15 | 5 | 32 | 568 | 31.9 | 75 | 0.55 | 10.5 | 52 |
| | L | 29 | 3 | 12 | 518 | 70.7 | 90.6 | 0.75 | 6.3 | 60 |
| 8 | R | 16 | 5 | 18 | 625 | 47.0 | 79.1 | 0.51 | 10.4 | 48 |
| | L | 21 | 4 | 14 | 656 | 60 | 84 | 0.72 | 8.3 | 70.0 |
| 9 | R | 28 | 5 | 17 | 612 | 62.2 | 84.8 | 0.76 | 9.9 | 65.3 |
| | L | 29 | 4 | 11 | 626 | 72.5 | 87.8 | 0.78 | 7.9 | 52.7 |
| 10 | R* | - | - | - | - | - | - | - | - | - |
| | L | 22 | 6 | 30 | 487 | 42.3 | 78.5 | 0.58 | 12.5 | 51.8 |
| 11 | R | 68 | 32 | 10 | 2588 | 87.1 | 68 | 0.74 | 20.2 | 67.5 |
| | L* | - | - | - | - | - | - | - | - | - |
| 12 | R | 34 | 6 | 12 | 830 | 73.9 | 85.0 | 0.76 | 9.4 | 63.5 |
| | L | 20 | 6 | 33 | 767 | 37.7 | 76.9 | 0.52 | 9.4 | 45 |
| 13 | R | 5 | 3 | 40 | 578 | 11.1 | 62.5 | 0.33 | 6.3 | 42.0 |
| | L | 2 | 3 | 47 | 529 | 4.0 | 40 | 0.24 | 6.3 | 45.0 |
| 14 | R | 20 | 5 | 13 | 721 | 60.6 | 80.0 | 0.72 | 9.9 | 72.0 |
| | L | 26 | 5 | 11 | 592 | 70.2 | 83.8 | 0.78 | 9.9 | 64.6 |
| 15 | R | 21 | 5 | 12 | 488 | 63.6 | 80.7 | 0.69 | 10.5 | 48.5 |
| | L | 21 | 5 | 8 | 599 | 72.4 | 80.7 | 0.75 | 10.5 | 65.7 |
| 16 | R | 57 | 16 | 132 | 2240 | 30.1 | 78.0 | 0.54 | 8.5 | 57.8 |
| | L | 76 | 12 | 61 | 1735 | 55.4 | 86.3 | 0.66 | 7.5 | 56.4 |
| **Sum** | **-** | **1122** | **295** | **1282** | **34720** | - | - | - | - | - |
| **Mean** | **-** | **-** | **-** | **-** | **-** | **50.2** | **77.9** | **0.61** | **9.95** | **55.3** |

*These cases did not show a clear beta peak in the power spectrum and were therefore excluded from further analysis.
